# Structures of the honeybee GABA_A_ RDL receptor illuminate allosteric modulation

**DOI:** 10.1101/2025.03.24.644576

**Authors:** Tatiana Labouré, Mayank Prakash Pandey, Eleftherios Zarkadas, Céline Juillan-Binard, Delphine Baud, Jacques Neyton, Thierry Cens, Matthieu Rousset, François Dehez, Pierre Charnet, Hugues Nury

## Abstract

Insect GABA_A_ receptors are critical insecticide targets, with all 21st-century commercial compounds acting through a single allosteric membrane site. Here we describe three ligand binding sites and the associated receptor conformations for the honeybee RDL receptor, combining cryo-EM, electrophysiology and molecular dynamics. First the conservation of the GABA site explains the absence of insect-selective orthosteric compounds. Second, the classical modulation site, occupied here by abamectin, exists in a closed-pore conformation contrasting with ivermectin-bound states in other receptors. Third, using the fungal-derived Chrodrimamin B, we resolve an unanticipated membrane modulation pocket structurally analogous yet pharmacologically opposite to a neurosteroid site in mammalian receptors. Structures also reveal the existence of an intersubunit, conformation-dependent, PIP2 lipid site. We foresee our results to be the starting point for investigations on the physiological modulation of insect GABA_A_ receptors. The honeybee receptor structures may also foster the search for environmentally benign insecticides.

## Introduction

Insecticides kill pest insects that would, in the absence of treatment, affect the yield of a crop. Insecticides are a pillar of agriculture. They are also the subject of polarized debates opposing their contribution to feeding the world’s population to their negative ecological impact. Neurotoxic molecules represent a large share of insecticides^1,2^ and often target ion channels.

γ-aminobutyric acid (GABA) is the main inhibitory neurotransmitter both in mammals and insects, where it plays a role in locomotion control, olfactory learning, and regulation of sleep and aggression. There are 4 genes encoding for ionotropic GABA (GABA_A_) receptor subunits in the honey bee (compared to 17 in humans), which assemble in specific pentameric combinations to form functional receptors^3^. Only the RDL subunit can assemble into functional homomeric receptors, while the GRD, the LCCH3 and probably the 8916 subunits need to be part of heteromeric receptors to yield GABA-elicited currents^4,5^. The RDL homopentamer pharmacology generally recapitulates that of native receptors^6^. However, most of our knowledge on the molecular operation mechanism of GABA_A_ receptors - and more generally on receptors from that same superfamily named pentameric ligand-gated ion channels (pLGICs)-stems from experiments performed with mammalian receptors^7,8^.

GABA_A_ receptors, from pest and beneficial species alike, are the targets of several classes of currently used insecticides as well as several older compounds^9,10^, all of them small molecules. Several generations of pore blockers (e.g. dieldrin in the 1950-60s, fipronil in the 1980-90s) found success before being banned due to toxicity to mammals or beneficial insects. New allosteric inhibitors emerged in the 2000s. Those include broflanilide, fluralaner^11,12^, fluxametamide^13^ and isocycloceram^14^. All those recent compounds bind to a membrane site overlapping with the site of the avermectins^15,16^, which are natural compounds used in crop protection (e.g. for potatoes and in horticulture)^17^ beyond their well-known anti-helminthic role in human health.

The use of proteins as clinical drugs was a revolution in the treatment of many human diseases. An equivalent revolution is yet to come for neurotoxic insecticides, with only one toxin-derived proteinic insecticide named Spear, on the market to date, targeting nicotinic acetylcholine receptors^18^. Proteinic insecticides offer advantages over small molecules: enhanced specificity and absence of long-term pollution. The manufacture, storage and above all, the delivery of protein-based neurotoxic insecticides, however, are enormous industrial obstacles. In any case, structures of pest targets and beneficial insect counter-targets will inform that revolution.

With the ultimate motivation of providing molecular information useful for the design of more specific insecticides, we report here cryoEM imaging of the *Apis mellifera* (the common honeybee) homomeric RDL receptor, in complex with GABA, alone, in complex with chrodrimanine B (ChroB) and in complex with abamectine (ABA). The set of four experimental structures, complemented by apt molecular dynamics (MD) and electrophysiology, constitutes a solid basis to understand the determinants of small molecule insecticide action. The results also point to unexpected avenues for physiological modulation by lipids or hydrophobic compounds.

## Results and discussion

### Agonist binding in the orthosteric site, in a possibly desensitized conformation

Among the 6 homologues tested, we identified the common honeybee RDL receptor as the most biochemically well-behaved (Fig. S1). The optimized sample for cryoEM harbors a fluorescent protein in lieu of the intracellular domain (Fig. S2) and is on-column reconstituted in a nanodisc delimited by a circularized, solubility-enhanced scaffold protein^19^. The RDL receptor has a typical pLGIC architecture with 5 subunits distributed symmetrically around the ion pore. In the agonist-bound structure, GABA occupy the 5 equivalent orthosteric sites located in clefts between subunits (Fig. 1). The binding pose of GABA is similar to that in the human receptors (Fig. 1c, S3). The neurotransmitter molecule lies within a cage of aromatic residues (Y239 from loop C, F191 from loop B, Y94 from loop D) that provide cation-π, anion-π, and hydrophobic interactions. The GABA amine also establishes interactions with E189 and the main chain carbonyls of S190 and F191, while the carboxylic acid contacts R96 and S161. Mutations of side chains in contact with GABA resulted in receptors that do not respond to the agonist (R96N, E189G and Y239A) or to a partial loss-of-function (F191A, EC_50_ 1 mM compared to 10 µM for the wild-type receptor, Table S1), in line with previously results observed for the *Drosophila* receptor^20^. An arginine from the loop E (R151) stabilizes loop C in a capped position that tightly covers GABA, reminiscent of the human ρ1 receptor^21^. The R151A mutant also yielded a drastic loss of function (EC_50_ in the mM range, Fig. 1g). In MD simulations featuring optimized parameters for cation-π interactions (see Methods), GABA molecules remain bound in the wild-type receptor but sometimes diffuse away in mutants, in particular in Y239A. Concomitantly, GABAs fluctuations are larger, and the loop C is more flexible for mutants (Fig. 1i).

**Fig. 1.**
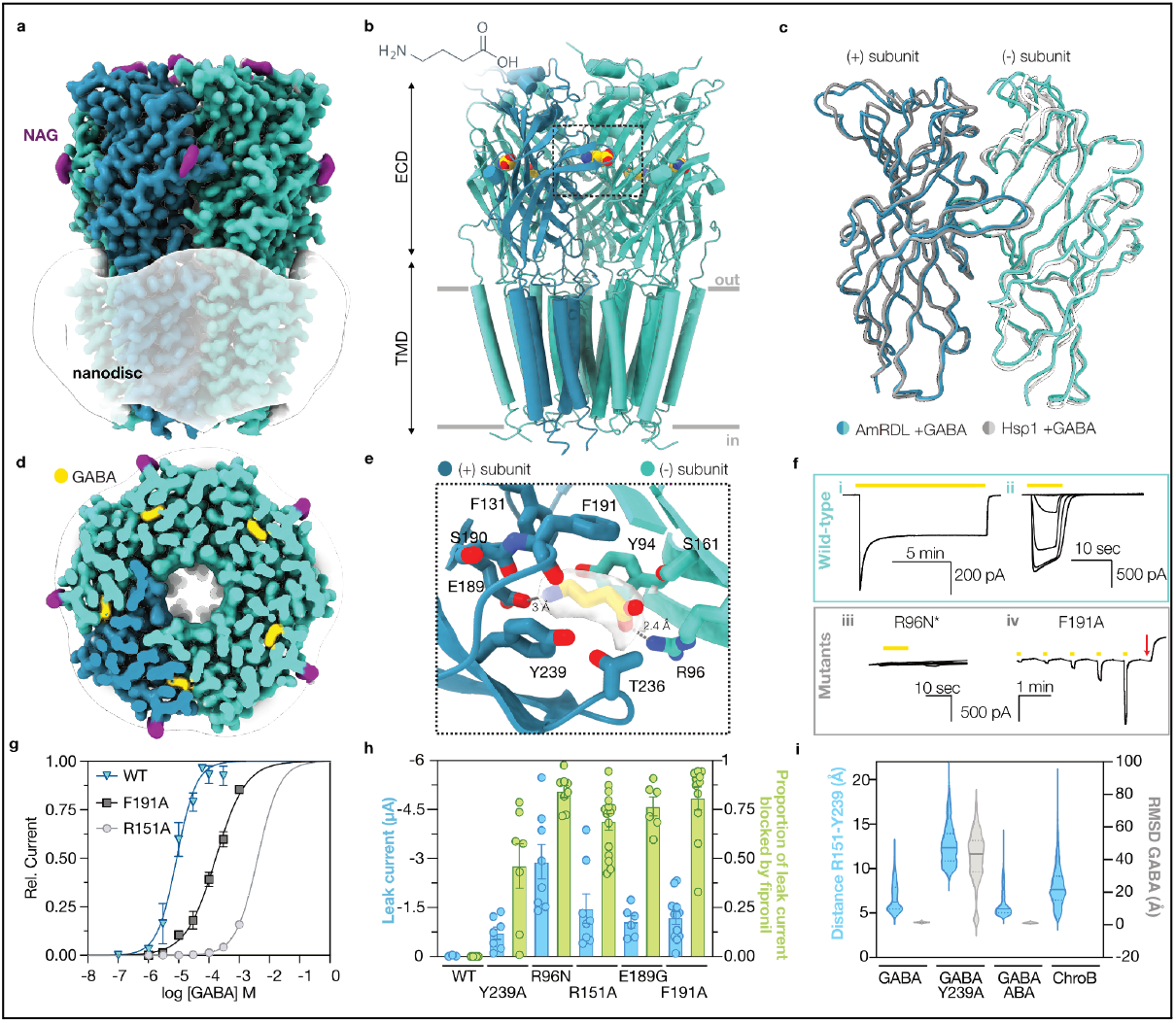
Binding of GABA in the orthosteric site. **a.** CryoEM maps of the AmRDL receptor in the presence of GABA, side view and **d**. cut-through top views. **b.** Structure of the receptor with bound GABA (yellow), side view. **c.** Overlay of two ECDs of the GABA-bound human ρ1 receptor (gray, 8OP9) and of the AmRDL receptor (blue). **e.** Orthosteric site close-up, with GABA and its surrounding residues shown as sticks. The density corresponding to GABA is shown. **f.** Representative current traces upon application of GABA (yellow bars) to oocytes expressing the WT receptor (i, ii) or mutants (iii, iv). Wild-type: (i) response to a long application of 100 μM GABA and (ii) responses to short GABA applications at 1, 3, 10, 30, 100, and 300 μM. Mutants: (iii) same as ii, absence of response for the R96N mutant. *The E189G and Y239A mutants have similar phenotypes. (iv) same as ii, followed by the application of 100 μM fipronil (red arrow) for the F191A mutant. **g.** GABA concentration response relationships for the WT receptor, the F191A and the R151A mutants. Means ± SEM data (n ≥ to 4) are plotted. **h.** Bar graphs showing the initial leak current (blue, μA) and the proportion of this leak current blocked after a 10-second application of 100 μM fipronil. Individual data points are plotted, together with mean and SEM (n > 4). **i.** Stability of the loop C (R151-Y239 distance, blue) and of GABA molecules (RMSD, gray) during MD simulations. Full line and dotted lines correspond to median and 1st/3rd quartile respectively.

Many of the orthosteric site mutations provoke large leak currents in oocytes, up to the µA range, that are partially inhibited by the specific pore blocker fipronil, indicative of spontaneous activation (Fig. 1h). Our findings thus indicate that site residues play a role both in binding and gating. In mammals, the β3 GABA_A_ receptor subunit is associated with spontaneous activation and plays a role in tonic inhibition. Residues responsible for the spontaneous activity are located within the loop F of the orthosteric site^22^. It is tantalizing to note that a picrotoxin-sensitive tonic inhibition was observed in honeybee Kenyon cells^23^. Future research might try to identify if the RDL or other GABA_A_ receptor subunits are involved in that tonic inhibition.

Superimposition to the agonist-bound ρ1 receptor (8OP9) shows clear similarities in the ECD conformation and clear differences in the TMD (Fig. 1c, S4). Those differences are not immediately apparent when comparing the pore profiles, which are globally similar with a constriction at its intracellular end, next to P2’ (P282) (Fig. 2c). However, the organization of the M2 helices strongly differs. Not only does the bundle of M2s not overlap well when the entire receptors are superimposed, but the interactions of each M2 with its neighbors are different, as observed when only one M2 is used for the superimposition (Fig. S4b).

**Fig. 2.**
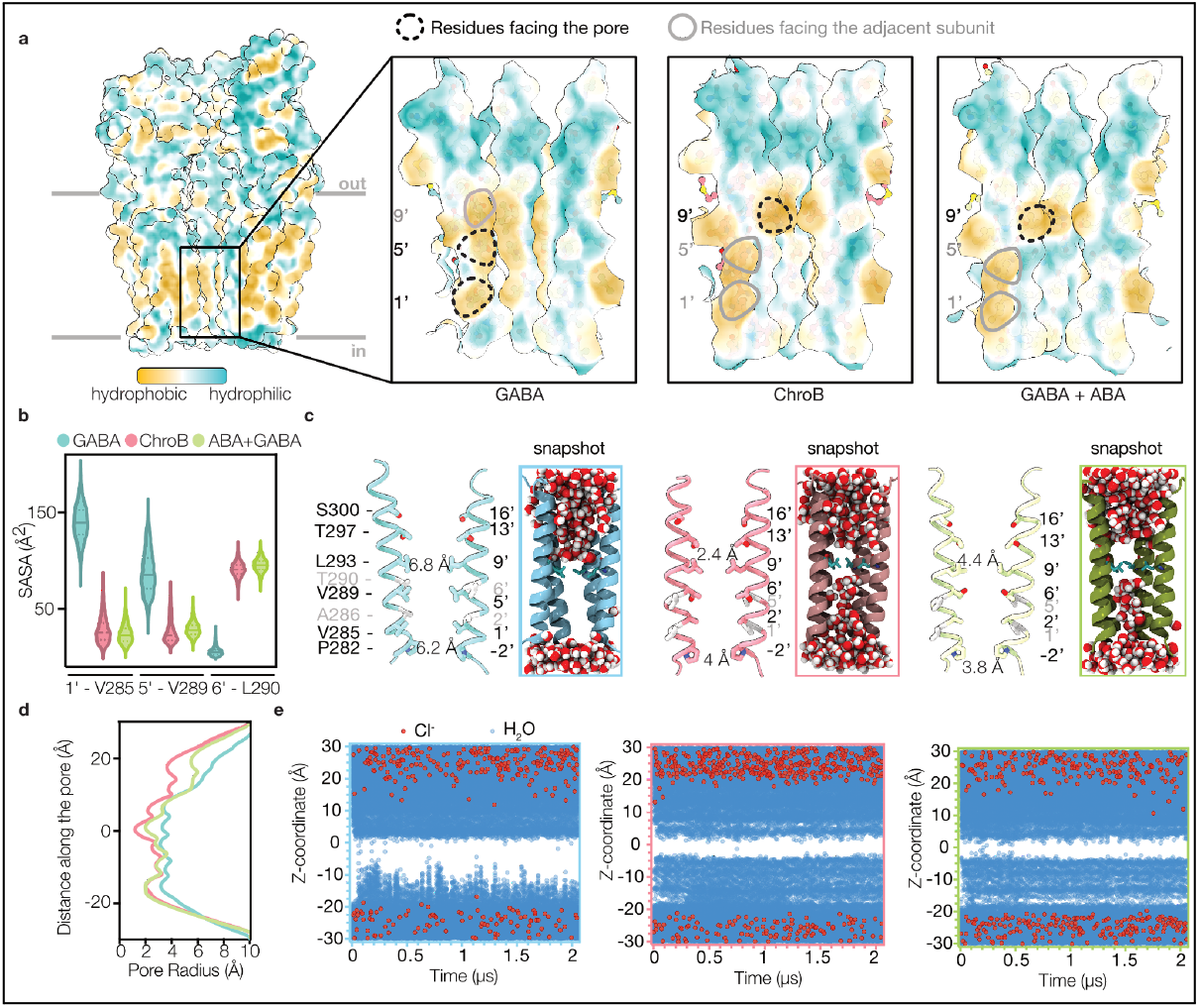
Pore geometry and permeation. **a**. Surface representation of the AmRDL receptor, showing the hydrophobic score with the two foreground subunits removed to reveal the pore. A zoomed-in view of the pore is included. **b**. Violin plot showing the solvent-accessible surface area (SASA) of M2 residues lining the pore during molecular dynamics (MD). **c**. M2 helices from two opposing subunits, with helices colored according to structure: GABA-bound (blue), ChroB-bound (pink), and ABA+GABA-bound (green). Numbers indicate pore diameters in Å at different positions, measured using CHAP. Snapshots of hydrated pores are shown. Red balls correspond to water molecules. **d**. Pore profile traces comparing the GABA-bound, ChroB-bound, and ABA+GABA-bound states of AmRDL. Distance 0 corresponding to Leu 9’ **e**. MD simulations showing pore hydration of the AmRDL structures, from the left to the right, GABA, ChroB and GABA-ABA-bound. Water molecules are represented as blue circles and Chloride anions as red circles. Z-coordinate 0 corresponds to Leu293 (9’).

Concomitantly the pore-exposed residues partially differ in the two structures. In both cases (and as in most pLGIC structures) side chains from residues 9’,13’,16’ face the extracellular moiety of the pore. At the intracellular end of the pore, however, residues 1’ and 5’ (V285 and V289) face the pore in the GABA-bound RDL receptor structure (Fig. 2a). The pore exposition of 1’ and 5’ appears stable in MD runs as evidenced by the residues’ solvent-accessible surface areas (Fig. 2b). Better than static pore profiles, MD simulations provide insights on permeation. We observe de-wetted pores in all replicas, as expected for a structure obtained in desensitizing conditions, in a region spanning 1’ to 9’ (Fig. 2e).

Such a pore organization is unusual. There is ample pLGIC literature that agrees on two rings of polar residues at 2’ and 6’ exposing side chains to the bottom pore lumen^7,8^. Also, the phenotype of the resistance mutant A2’S for pore-blocking insecticides^24^ has a straightforward explanation in the framework of 2’ facing the pore. The data can thus be interpreted in two ways. On the one hand, the unusual pore geometry could be a unique physiological feature of the desensitized conformation of the AmRDL receptor. On the other hand, the pore geometry could be an artefactual consequence of experimental conditions (for instance the lipidic composition of the nanodiscs) not reflecting a physiological state.

We next recorded a dataset in the absence of any ligand, aiming to capture the resting state. In the apo structure, the loop C is uncapped, with an outward motion of ~5Å at its tip (Fig. S5). The loop C density is less well-resolved, indicative of an increased flexibility in the absence of a ligand. The orthosteric site becomes fully accessible to the solvent and is empty. At the TMD level, the map indicates a high degree of flexibility (Fig. S5a). To our surprise, the global conformation does not present the hallmarks of a resting state conformation: we had expected to see a clear ECD/ECD interface reorganization, and a drastic closure of the activation gate around the 9’ position of the pore, but instead observed only limited motions. Among the structures determined here, the apo receptor resembles the GABA-bound conformation more than the other ones (the differences are discussed in Fig. S5). We wondered if adding an antagonist might shift the equilibrium of conformations and yield a different closed-pore state.

### Chrodrimanin B binds to a membrane site, next to a PIP2 site, and stabilizes an inhibited conformation

While the mammalian receptor antagonists bicuculline and gabazine are not potent RDL antagonists, a fungal compound named ChroB has been described as a strong competitive antagonist. ChroB kills silkworm larvae and inhibits their RDL receptors at nanomolar concentrations, while being about 1000-fold less potent at the human α1β2γ2 receptor^25,26^. Because it decreases the binding of radioligands targeting the orthosteric site but not the pore, ChroB was described as a competitive antagonist and modeled in the neurotransmitter site^27^. However, in cryoEM datasets obtained in the presence of 10 µM ChroB, the 3D reconstructions feature an uncapped loop C over an empty orthosteric site (Fig. 3). Instead, ChroB binds to a mostly hydrophobic site in the inner membrane leaflet, where a banana-shaped density is present in between subunits. In simulations with ChroB, the ligands establish stable hydrophobic interactions with A334, L331 and F327. The ChroB O7 oxygen atoms also form H-bonds with T335 and with Y338 (Fig. 3g), the latter alternating between ChroB and a PIP_2_ lipid (see below). We observe water molecules penetrating in the bilayer zone and bridging ChroB O8 oxygens with W275 indole’s nitrogen (Fig. 3a,c). Mutations of nearby residues profoundly impact the effect of ChroB: inhibition is abolished for A334L, T335L, W275L and Y338L. ChroB strikingly turns into a strong positive allosteric modulator (PAM) for Y338A, increasing the GABA-elicited currents by ~15-fold at the EC_30_ (Fig. 3d-f). The combined structural, functional and simulation data confidently identify a novel allosteric site for RDL receptors, the targeting of which may foster the development of insecticides with a new mode-of-action.

**Fig. 3.**
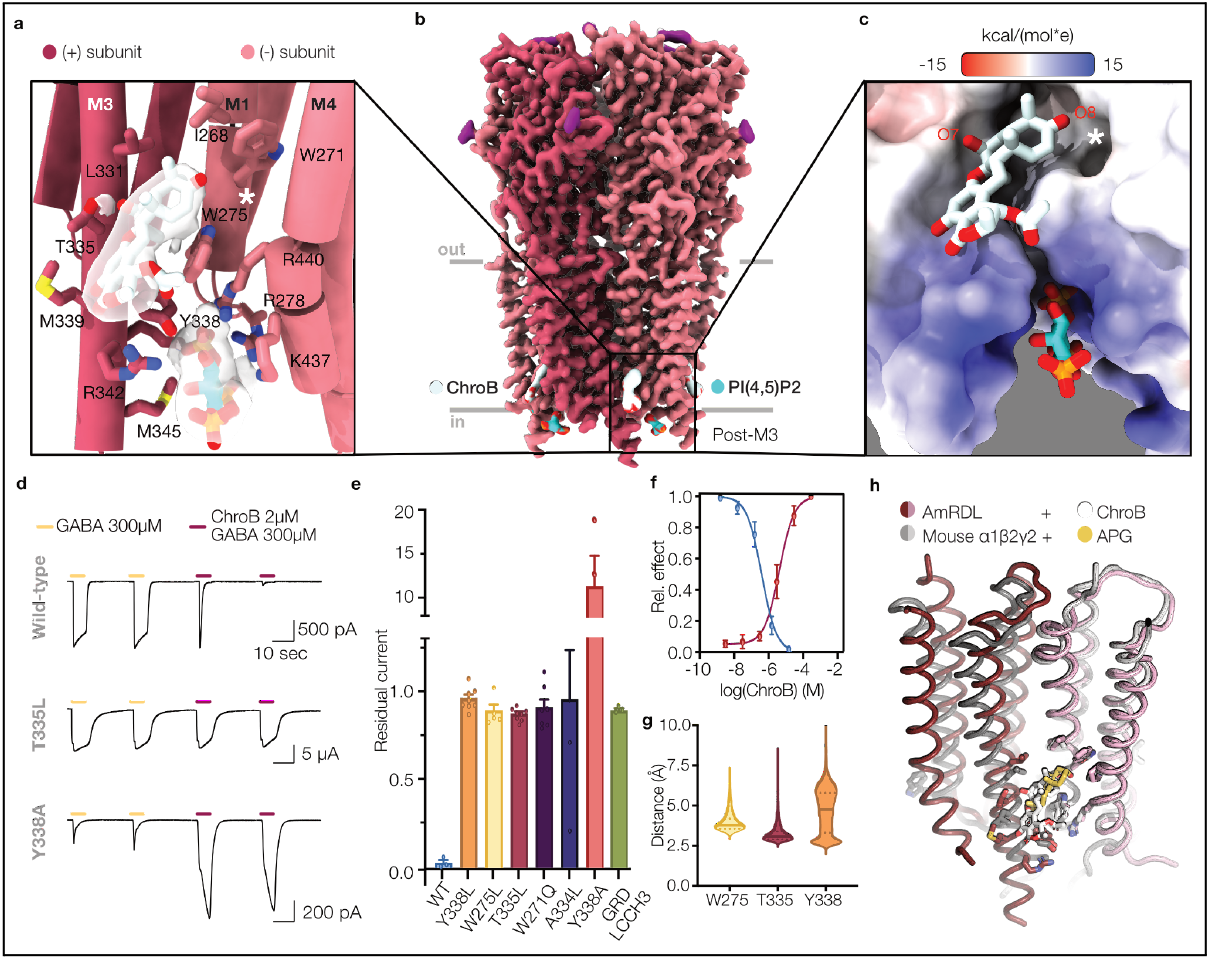
ChroB binds to an intersubunit membrane site, close to the cytoplasm, next to a PIP_2_ site. **a**. Close-up view of the ChroB (white) and PIP_2_ (cyan) binding pockets, with ligands and lining residues shown as sticks. The densities around the ligands are shown. The white star indicates the presence of a water molecule observed in molecular dynamics (MD) simulations. **b**. Cryo-EM map of the AmRDL receptor with bound ChroB and PIP_2_. **c**. ChroB and PIP, are represented as sticks on the receptor surface, colored according to electrostatic potential. PIP_2_, binds to a highly electropositive pocket. **d**. Representative current traces upon application of 300 μM GABA or 300 μM GABA + 2 μM ChroB to oocytes expressing WT or mutant (T335A and Y338A) receptors. **e**. Residual GABA-elicited currents after two applications of 300 μM GABA + 2 μM ChroB. Individual data points are plotted, together with mean and SEM (n ≥ to 3) **f**. ChroB dose-response dependencies for the WT receptor (relative inhibition, blue) and the Y338A mutant (potentiation relative to the effect at 200μM, pink) **g**. Distances between ChroB and key binding pocket residues during MD simulations. **h**. Overlays of two TMDs of the ChroB-bound AmRDL receptor with the allopregnanolone(APG)-bound murine α1β2γ2 receptor (8FOI).

Interestingly ChroB occupies a site equivalent to the PAM neurosteroid site in human GABA_A_ receptors^28^. Brain neurosteroids are physiological modulators with a therapeutic potential, as illustrated by recent FDA approvals of ganaxolone, allopregnanolone and zuranolone. Superimpositions of the ChroB-bound RDL receptor to neurosteroid-bound receptors show similarities in the ligands orientation and coordination: for instance, W275, Y337 and L331 are conserved (Fig. 3h, S6). The different subunit/subunit arrangement, clearly apparent when looking at the principal subunit upper M2 and M3 (Fig. S6e), nevertheless echoes the opposite action of PAM neurosteroids and NAM ChroB. The existence of an innate insect molecule acting through the ChroB site is a far-fetched yet fascinating hypothesis for future studies. Here, we conducted a limited exploratory experiment, assessing the impact of a chemically related ecdysone hormone on the receptor surface expression. In the overexpression conditions of the baculovirus-infected Sf21 cells, 2-hydroxy-ecdysone (20-HE), the major molting hormone, increases the amount of RDL receptor present at the plasma membrane (Fig. S6d).

In the ChroB reconstruction, we noticed an extra density featuring three protrusions, surrounded by positively charged residues (R278, K341, R342, K437, K440 - not all side chain densities are well-defined). It lies next to ChroB at the intracellular end of M3 and M4. We modeled there the inositol-4,5-triphosphate head-group of PIP_2_ (Fig 3b, c and S7). The entire PIP_2_ lipids, modeled in MD simulations, exhibited tightly bound head-groups, stabilized by a network of salt bridges (Fig. S7), and flexible acyl tails. Of note, a chloride binding site was found in simulations in between PIP_2_, R278 and the pore exit mouth. A PIP_2_ density was present in the GABA+ABA-bound cryoEM map - at lower contouring-but is absent in the GABA-bound reconstruction, in line with the conformation being drastically different in that area (Fig. S7, see below). Most of the basic side chains coordinating PIP_2_ are conserved and have an equivalent role in the human GABA_A_ receptor^29^. However, those from the M1-M2 loop and M4 are oriented towards the opposite side of the M4 helix. Evolution thus yielded structurally distinct sites: an insect inter-subunit PIP_2_ site instead of the human intra-subunit site, while conserving sequences (Fig. S7e). PIP_2_ participates in the trafficking of mammalian GABA_A_ receptors. In addition, receptors diffuse from cholesterol-rich micro-environment to PIP_2_-rich ones upon exposure to GABA, with an impact on endocytosis^30^. Future studies should delineate the role(s) of PIP_2_ in the function or localization of insect receptors.

The pore adopts the typical pLGIC geometry closed conformation, with side chains of residues −2’, 2’, 6’, 9’, 13’, 16’ facing the pore lumen. It features a constriction at the level of L9’ side chains where the pore is de-wetted in simulations (Fig. 2f). The dewetted zone is restricted to the vicinity of 9’, in contrast with simulations of the GABA-bound structure where it spans a thicker zone between. In oocytes injected with the L9’S mutant, we observed strong leak currents that could be blocked by fipronil (data not shown), corresponding to spontaneous activation, in agreement with previous reports^31,32^ imparting a crucial role of this residue in gating. Furthermore, the ChroB-bound structure resembles more the apo and inhibited conformations of the human ρ1 receptor (8OQ6 and 8OQ7) than the agonist-bound one (8OP9). Altogether, the data are fully consistent with the ChroB-bound structure representing an inhibited state.

### Abamectin binds to the interface between subunits, in the TMD

ABA, a macrocyclic lactone, is the natural precursor of the anti-parasitic drug ivermectin (IVM). Beyond its nematicide activity, ABA is extremely active against a range of arthropods^17,33^ and it inhibits the honeybee RDL receptor with an IC_50_ of 8 nM^16^. However, the effects of ABA are multifaceted. In the housefly, ABA is an agonist and a positive allosteric modulator at low GABA concentrations but becomes a negative allosteric modulator at high GABA concentrations^34^. ABA also targets the GluCl receptor of arthropods, which is often considered its main physiological target, based on the existence of resistant GluCl mutants.

We obtained cryoEM data in the presence of 3mM GABA and 1µM ABA. The former binds to the orthosteric sites. The latter binds tightly to mostly hydrophobic inter-subunit membrane cavities located just below the M2-M3 loops (Fig. 4). The cyclohexene moiety of ABA penetrates deep in between M3 from the principal subunit and M1 from the complementary subunit, to come in contact with M2 at M299 (M15’). The surrounding residues establish stable contacts with ABA during MD simulations (e.g. I257, G320, Fig. 4g). By contrast, the contacts are more labile with the spiroketal moiety, which lies at the mouth of the cavity, and with the di-saccharide moiety, indicative of the flexibility of these chemical groups. In MD trajectories, the removal of ABA led to a moderate constriction of the cavity, which becomes similar to that existing in the GABA-only structure, while the global conformation remains close to the experimental GABA-ABA structure at the time scale of the simulations. In both the GABA-only and the GABA-ABA without ABA runs, lipid tails frequently visit the empty cavities.

**Fig. 4.**
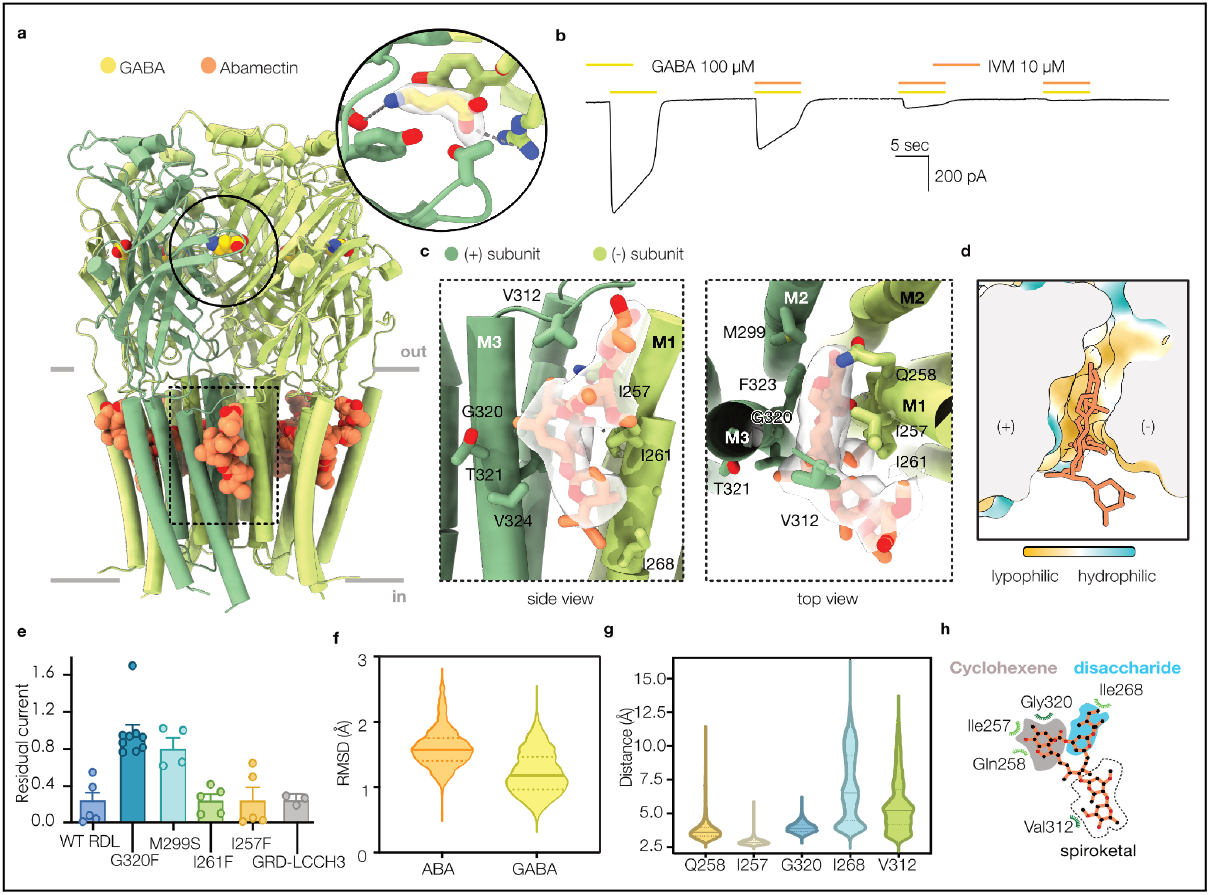
Binding of Abamectin in membrane site, GABA-ABA-bound receptor conformation. **a**.Structure of the AmRDL receptor with bound GABA (yellow) and Abamectin (ABA, orange), viewed from the membrane plane. Close-up of the GABA-occupied orthosteric site. **b**. Representative current traces from oocytes expressing the AmRDL receptor, showing responses to 100 μM GABA with or without 10 μM Ivermectin (IVM). The sole difference between IVM, used in all functional experiments, and ABA is that ABA bears an olefin bound between C22 and C23, while ivermectin is the 22,23-dihydro derivative. **c**. Side and top views of the ABA binding pocket, with the ligand and neighboring side chains shown as sticks. **d**.Surface representation of the ABA pocket, top view, colored by hydrophobicity. **e**.Inhibition by IVM: residual GABA-elicited currents after three applications of 100 μM GABA + 10 μM IVM in oocytes expressing WT or mutant AmRDL receptors and the heteromeric AmGRD-LCCH3 receptor. The two isoleucine mutants correspond to RDL-LCCH3/GRD substitutions **f**. RMSD of both GABA and ABA molecules during MD simulations. **g**. Distances between ABA and binding pocket residues (schematized in **h**) during MD simulations.

We designed mutations to sterically perturb different parts of the cavity. The biggest effect was obtained for the G320F mutant, which almost abolished the negative modulation, but was also 50-fold less sensitive to GABA (Fig. 4d,e). This glycine residue is conserved across invertebrates and harbors the most common resistance mutations found in the fields for ABA and a host of other compounds binding to the same pocket. At the molecular level, it was shown that G320A and G320M fully abolished the ABA inhibitory effect on the housefly RDL receptor, with G320A strongly increasing the PAM effect of the compound^34^.

IVM has been previously observed in the same site in the *C. elegans* GluCl receptor^35^ and glycine receptors^36,37^, the latter being affected at much higher concentrations by the drug. Deep similarities are seen in the binding pauses of IVM in GluCl and ABA in the RDL receptor (Fig. S8). In particular, the ligand lies on M3 in a very similar way, due to the conservation of ‘bottom wall’ F327 and the ‘left wall’ G320. Mammalian glycine receptors have an alanine at that position, resulting in IVM being displaced ~1.5Å away from M3. ABA slides ~1 Å deeper against M1 in the RDL receptor than IVM does in GluCl. This modestly deeper penetration correlates with the position of M2. The interaction at the level of residue 15’ from M2 is one of the few non-conserved contacts, with the RDL receptor bearing a methionine and GluCl a shorter, more polar, serine. Interestingly, the M15’S mutant has the second strongest effect in the series tested. The side chain at 15’ has also been identified as the largest determinant of the level ivermectin activation in the different isoforms of GluCl from the sea louse^38^.

A major difference between the ABA-bound structure and previously determined IVM-bound structures is the closure of the pore at the 9’ level (Fig. 2). In GluCl and in glycine receptors, IVM binding is associated with a relatively large diameter at the level of 9’, compatible with the passage of ions^35–37,39^, while the level of constriction at −2’ can vary. In stark contrast, we observe a closed hydrophobic gate with the side chains of L9’ oriented towards the pore. In simulations starting with the ABA-bound structure, the pore remains un-wetted at the 9’ level, both with and without ABA, underlining the metastability of the pore conformation (Fig. 2g). Given the dual effect of avermectins, with an inhibitory action at high neurotransmitter concentration, one does expect a closed-pore conformation at the saturating concentrations used for both ligands in the cryoEM experiments. With its constriction at 9’, our ABA-bound structure defines a new conformational template, with a small-but-significant cavity geometry difference, that would be useful for the design of compounds targeting the same site. This would be key for compounds derived from current insecticides that bind to this site, such as fluxametamine, isocycloseram or fluralaner.

### Comparison between structures, assignment to states

We next compared the three different conformations at several structural levels (Fig. 5). There is very little intra-domain tertiary motion for the ECD: each ECD moves as a rigid body except at the level of the loop C. In the TMD, the tertiary deformation is limited to M2 tilting relative to the other helices. The transition thus essentially results from subunit/subunit reorganization, which we next describe from the extracellular side down to the intracellular end of the pore. First, comparing the ChroB structure with the GABA-bound ones at the ECD level, there is a ~2° degree rotation of a subunit ECD versus the next (Fig. 5c) and a compaction of the subunit-subunit interface. At the level of the pentamer, this quaternary reorganization results in a counter-clockwise rotation and the spread of the β6-β7 cys-loop, the β1-β2 and β8-β9 loops, away from the pore axis. Second, at the interfacial level between the ECD and the TMD, the M2-M3 loop slides relative to the β1-β2 loop (with P307 moving past V85, Fig. 5b), a feature associated with activation^40^. Third, comparing the ChroB structure with the GABA-ABA one at the TMD level, there is a rotation of a subunit TMD versus the next that corresponds to a widening of the extracellular moiety of the M2 pore helices. Fourth, comparing the GABA-ABA structure with the GABA structure, the upper TMD remains similar while the lower TMD is re-arranged. Large amplitude motions take place at the intracellular ends of helices. We identified residues, which pair distances function as a signature of either the GABA conformation (T281 far from L276; A280 close to G337) or of the inhibitor-bound conformations (T281 close to L276; A280 far from G337) (Fig. 5h-i). Those distances are stable during simulations of the different states.

**Fig. 5.**
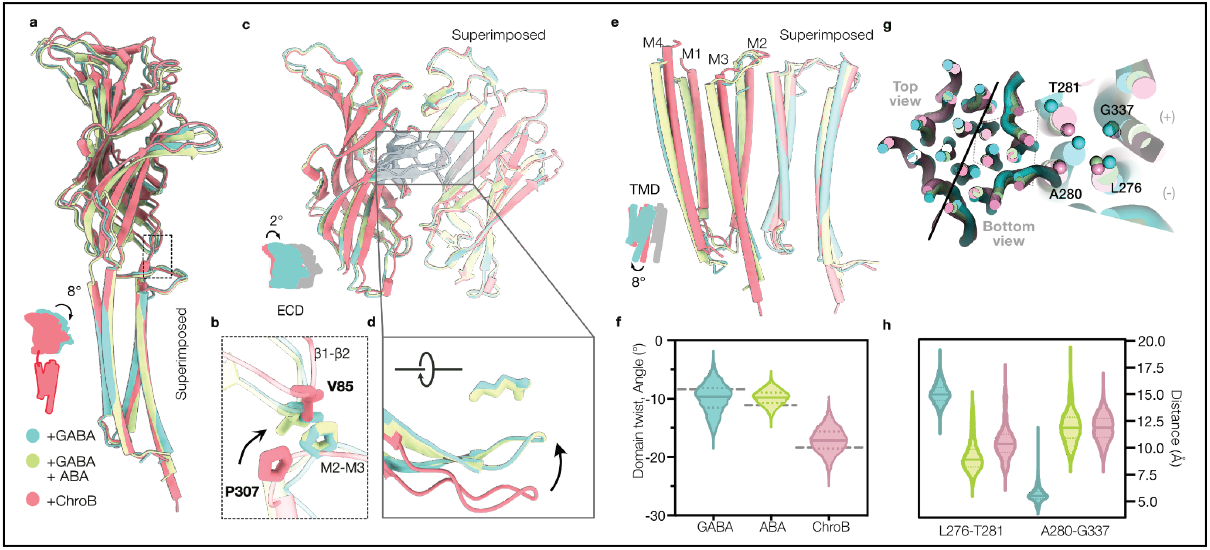
Comparison of the conformations obtained in the presence of GABA, of GABA+ABA or in presence of ChroB. **a**. Overlay of subunits bound to GABA (blue), GABA+ABA (green), and ChroB (pink), viewed from the membrane plane, showing the ECD/TMD rearrangement. **b**. Close-up views of interactions between the β1-β2 and M2-M3 interfacial loops. **c**. Overlay of two adjacent ECDs, highlighting the rigid body tilting of one relative to the other upon GABA binding. **d**. Close-up views of loop Cs highlighting capping. **e**. Overlay of two adjacent TMDs. **f**. Domain twists measured during molecular dynamics (MD) simulation. The dashed lines represent the CryoEM structure value. **g**. Overlays of the TMDs, viewed from the top and from the bottom, with a global TMD superimposition. Close-up on pairs of residues of the lower TMD (L276-T281 and A280-G337) reflecting the large local rearrangement. **h**. Violin plot of distances between the residues highlighted in (h) during MD simulations.

The unique conformation of the M2s in the GABA structure can be described as an iris-like transition. The motion is congruous with a stark increase in the tilt of the M2 helices relative to the pore axis, and to a clockwise rotation of the lower M2 (seen from the extracellular side). While in the inhibitor-bound conformations, one M2 mainly contacts the neighboring principal subunit, interactions are switched to the complementary subunit in the GABA-bound structure.

Ever since the very first *Torpedo* nAChR structures, the correspondence between experimental structures and functional states has been established with varying degrees of confidence. This remains true here. With good confidence, we assign the ChroB bound structure to an inhibited state, on the basis of its closed pore and its global conformation being similar enough to many previously determined inhibited structures. With reasonable confidence, we believe the GABA+ABA-bound structure represents a new template, different from previously determined ivermectin-bound structures, for the state stabilized by a host of insecticides. In addition, with its agonist-bound ECD and its 9’ closed-pored TMD, the GABA+ABA-bound structure seems close to pre-open states. Finally, we provisionally assign the GABA-bound state to a desensitized state with a unique arrangement of M2 helices. Our findings point to several possible avenues for future research aimed at validating or disproving those assignments: looking at the influence of lipid composition or membrane mimetic type or using conformation specific crosslinking.

## Conclusion and outlook

We provide a structural framework that reveals ligand binding sites and the conformational landscape of the RDL receptor from a beneficial insect, the honeybee. We believe our work is opening avenues that go in different directions. For instance, future investigations could reveal the role of PIP_2_, structure-function studies may shed light on the dual action of avermectins and single-channel work could explain the mechanism behind spontaneous openings. Furthermore, GABA, ABA and ChroB all bind at subunit interfaces and mutating the binding pockets could help define insect heteromeric receptor composition. The pharmacology of the ChroB site, which is not currently targeted by any commercial insecticide, will certainly be explored and might lead to the discovery of insecticides with a novel mode-of-action. Of course, the honeybee RDL should not be targeted by insecticides. On the contrary, it must be a counter-target. A long-term exciting direction will be the development of protein modulators with insecticidal properties. Protein design is undergoing a Cambrian explosion and its possibilities are quickly expanding. For the well-established targets of the pLGIC family, the reported work might be an initial step in enabling the design (by providing templates) and the selection (by providing the recipe to produce the pure protein for negative selection) of future environmentally benign neurotoxic insecticides.

## Methods

### Materials

Chemicals were purchased from Sigma unless stated otherwise. Lipid and detergent were purchased from Avanti. Chrodrimanin B was purchased by BioAustralis Fine Chemicals.

### AmRDL receptor production

Two constructs were generated, corresponding either to the full-length AmRDL receptor (Genbank KJ485710, dataset with GABA+ABA) or to a truncated receptor lacking the intrinsically disordered domain (residues 334 to 429, apo, with GABA, with ChroB datasets), which was replaced by the GFP. Both constructs harbored a C-ter TEV site and a rhodopsin-1D4 tag (TETSQVAPA). The constructs were inserted in the pFastBac vector using In-Fusion cloning (Takara Bio). The receptor was expressed in baculovirus-infected insect cells. Briefly, *Spodoptera frugiperda* 21 cells (Sf21, Invitrogen) were maintained at a density of 0.7×10^6^ per mL in serum free Sf-900 III SFM medium at 27°C under 90 rpm agitation. Baculoviruses were generated by transfecting 10^6^ Sf21 cells with 5 μg of purified bacmid and by harvesting the supernatant, called V_0_, 72h later. The V_0_ was then amplified to infect a large volume of cells (typically 2-4 L at 0.9×10^6^ cells /mL infected at 1/100 V/V). Cells were harvested 72 h post-infection.

### Immunofluorescence microscopy

Sf21 cells were infected with AmRDL baculovirus in the presence or absence of 40 μM 20-hydroxyecdysone (Santa Cruz biotechnology) and incubated for 72 hours. Cells were fixed with 4% paraformaldehyde (PFA) diluted in PBS during 20 min followed by 20 min non-specific site saturation step in PBS-BSA 2% (w/v). The cells were incubated with an anti-histag antibody (Thermo Fisher) for 60 min and nucleic acids were stained with DAPI during 10 min. Mounting of coverslips was done using Mowiol medium. The images were acquired with a confocal microscope (S-M4D, Olympus GATACA, and Andor) on a motorized IX81 stand, equipped with two EMCCD cameras (Andor iXon ultra).

### AmRDL receptor purification and nanodisc reconstitution

Sf21 cells were resuspended in buffer A (20 mM Tris-HCl pH 7.4, 300 mM NaCl) supplemented with a protease inhibitor cocktail and mechanically lysed. Membranes were recovered by ultracentrifugation at 100’000 g for 1 hour and then resuspended in buffer A, supplemented with a protease inhibitor cocktail, 20 mM MgCl_2_, ATP, and DNase, followed by mechanical homogenization. 8 g of membranes were solubilized using 1% n-dodecyl-β-D-maltoside (DDM) and 0.1% cholesterol hemisuccinate (CHS) under gentle agitation for 1.5 hours. The soluble fraction was collected after ultracentrifugation (100’000 g for 0.5 hour) and incubated with 500 μL of 1D4 affinity resin (CubeBiotech) overnight. The beads were recovered by centrifugation and washed with 10 column volumes (CV) of buffer A supplemented with 0.02% DDM, 0.002% CHS, and 0.04 mg/mL porcine brain lipid (BL, stock 5 mg/mL in 3% DDM).

To reduce purification time, the receptor was inserted into nanodiscs while attached to the affinity resin as described in ^29^. After washing, the resin was incubated with 0.4 mg/mL of circularized MSP1E3D1 and 0.4 mg/mL of BL for 0.75 hour under gentle agitation. To remove the detergent, the resin was incubated with two rounds of Bio-Beads for 0.75 hour each. The beads were recovered by centrifugation and washed with 10 column volumes of buffer A. The receptors were eluted with buffer B (20 mM Tris-HCl pH 7.4, 150 mM NaCl) supplemented with 500 μM Rho1D4 peptide in two rounds of 2 CV, for 1 hour each. The eluate, containing 0.05–0.1 mg/mL of protein, was concentrated on a 100 kDa cut-off concentrator to reach a concentration of 1 mg/mL, suitable for the preparation of a few cryoEM grids.

### CryoEM

Purified AmRDL receptor in nanodiscs was incubated for 0.5 hours with the megabody Mb^c7HopQ^NbF3 (Uchański et al., 2021) (targeting the scaffold protein) at a 3:1 megabody:receptor molar ratio. Grids were prepared by depositing 3.5 μL of the sample onto a glow-discharged (30 mA, 50 s) UltrAuFoil Au 300 R 1.2/1.3 grid, waiting 6 s, and blotting for 5 s at 8°C with 100% humidity using a Mark IV Vitrobot (Thermo Fisher Scientific), followed by plunge-freezing in liquid ethane. For insecticides datasets, the compound was added 5 min before freezing. The datasets were recorded either on a Titan Krios Cryo-TEM (Thermo Fisher Scientific) at the European Synchrotron Radiation Facility (ESRF) or on a Glacios Cryo-TEM (Thermo Fisher Scientific) at the Institut de Biologie Structurale (IBS), as detailed in Table S2.

Data were analyzed using CryoSPARC following a workflow illustrated in Fig. S9, except for particle picking, which was performed using crYOLO. Poor-quality micrographs were excluded through manual inspection, and picked particles were classified into 2D classes. Only 2D classes resembling pentameric assemblies were kept for further analysis. *Ab initio* reconstruction and heterogeneous refinement were used to curate particles in 3D. The final 3D reconstruction was obtained after several iterations of non-uniform refinement with C5 symmetry applied. An AlphaFold2 model was used as the initial atomic model and fit as a rigid body in the cryoEM map. The model was refined through several rounds of manual rebuilding using Coot (Emsley et al., 2010) and of real-space refinement using Phenix (Liebschner et al., 2019) and Isolde^41^. Maps post-treated with DeepEMhancer were used for visualization. Figures were prepared with PyMOL (Schrodinger) and ChimeraX (Pettersen et al., 2021). Pore diameters were calculated using CHAP (Trick et al., 2016). Root mean square deviations (r.m.s.d.) were calculated using ChimeraX.

### Molecular dynamics simulations

The experimental structures served as the starting points for the simulations. Lipid bilayer systems were constructed using CHARMM-GUI^42,43^, embedding each structure in a hydrated POPC (1-palmitoyl−2-oleoyl-sn-glycero-3-phosphocholine) bilayer containing approximately 378 POPC lipids. Ligand parameters were generated using CGenFF^44^. Parameters associated with five dihedral angles involved in the dihydropyran ring connected to the acetate ester of CroB exhibits relatively large CGenFF penalties. Therefore, these dihedrals were harmonically restrained to their experimental values during our simulations, All systems were solvated with TIP3P^45^ water and neutralized with 150 mM NaCl, resulting in systems with approximately 210,000 atoms and initial dimension of 120 x 120 x 150 Å^3^. Simulations were performed using NAMD-3.0^46^ with the CHARMM36m^47^ force field. Dedicated NBFIX parameters^48^ were used for cation-pi interactions between ligand and protein. Long-range electrostatic interactions were calculated using Particle mesh Ewald^49^. Temperature control was managed via Langevin dynamics at 300 K, with a damping coefficient of 1 ps^−1^. Pressure was regulated using a semi-isotropic Langevin piston^50^ at 1 atm. The energy of each system was first minimized for 3000 steps. Next, all molecular assays were equilibrated over 10 ns with harmonic restraints maintaining the backbone and the side chain atoms to their CryoEM positions. Subsequently, side chain restraints were removed, and the system underwent further equilibration for 40 ns restraining only backbone atoms. The latter restraints were gradually released over the next 5 ns. Finally, production simulations ran for 2 μs with timestep of 4 fs using hydrogen mass repartitioning (HMR)^51^. Three independent replicas were performed for each system. Visualization and trajectory analysis were conducted using VMD^52^. The domain twist was calculated as the dihedral angle formed by the center of mass (COM) of the ECD of a subunit, the COM of the ECD of all subunits, the COM of the TMD of all subunits and the COM of the TMD of a subunit^21^.

### Two-electrode voltage clamp electrophysiology

The cDNA encoding the AmRDL subunit (Genbank KJ485710) was cloned into the pBluescript-II cloning vector (Agilent Technologies, Inc.) and the TEVC mutants were generated using the NEBuilder HiFi assembly kit (NewEngland Biolabs, Ipswich, MA, USA) and oligonucleotide pairs designed using the NEbuilder® assembly tool (https://nebuilder.neb.com). Two fragments were amplified by PCR using the Herculase II fusion polymerase (Agilent) using AmRDL cDNA as a template. The fragments were cloned into the pcDNA3.1(+) vector, with the alfalfa mosaic virus sequence immediately before the start and the 3’ UTR sequence of the *Xenopus laevis* β-globin gene immediately after the stop codon. Mutation and sequence integrity were verified by sequencing the fragments on both strands. For *X. laevis* oocyte injections, cRNAs were prepared from linearised plasmids using the Mmessage Mmachine Transcription Kit (Thermo Fisher) according to the manufacturer’s instructions.

Oocyte preparation and injection were performed as previously described (Henry et al., 2020; Rousset et al., 2017; Cens et al., 2015, 2013), and were carried out in strict accordance with the recommendations and relevant guidelines of our institution. Surgery was performed under anesthesia, and efforts were made to minimize suffering. The experimental protocols were approved by the Direction Départementale des Services Vétérinaires (authorization N° C34.16). Oocytes were surgically extracted from anesthetized (MS-222, Sigma-Aldrich A5040, 0.2%) female Xenopus laevis and dissociated with collagenase 1A (Sigma-Aldrich C9891, 1 mg/ml) in a low calcium solution (mM: NaCl 82; KCl 2; MgCl2 1; HEPES 5; pH 7.2 with NaOH) for 1.5 hours. Groups of ~30 oocytes were pressure-injected (50–300 ms at 1 bar) with ~30–50 nl of RNA (or cDNA, 0.1–1 µg/µl) and incubated at 19°C in NDS solution (mM: NaCl 96; KCl 2; CaCl2 1.8; MgCl2 1; HEPES 5; Na-pyruvate 2.5; gentamicin 0.05; pH 7.2 with NaOH) for 1–3 days.

For two-electrode voltage-clamp recordings, pipettes (Harvard GC150T10, 0.2–1 MΩ) filled with 3 M KCl were used. The basic recording solution (mM: NaCl 96; KCl 3; CaCl2 0.5; MgCl2 1; HEPES 5; pH 7.2 with NaOH) was connected to the amplifier (Geneclamp 500B, Axon) via a virtual ground head stage and agar-KCl (3 M) bridges. Junction potentials were nulled with both pipettes in the bath before impalement, and the holding potential was −60 mV. Oocytes were perfused (1–2 ml/min) with recording solution, with 5–10 s pClamp-controlled (ver. 7.0, Axon) applications of GABA-containing solution (0.1–10,000 µM) every 45–90 s. Antagonists (ChroB, Fipronil, or Ivermectin) were applied alone or with GABA. Dose-response curves were generated by perfusing increasing GABA concentrations.

### Electrophysiology data analysis

Current trace analysis was performed with Clampfit (Axon Inst. Ver 10, Molec Dev.). GABA-induced increase in current was measured at the peak amplitude of the GABA-current relative to the current amplitude recorded before GABA perfusion. The dose-response curves for GABA were obtained by normalizing the currents obtained with the different concentrations of GABA to the highest current amplitude obtained (usually with the highest dose of GABA, i.e. 1 mM) and plotted against the log of the GABA concentration. All concentration–response curves were fitted with GraphPad Prism 8. The concentration response curves underlying Fig.1 and Fig.3 were fitted using log(agonist) vs. response – Variable slope (four parameters) with variable EC50. The following formula was used for fitting:

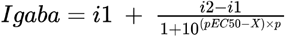

where Igaba is the current amplitude measured in the presence of different concentrations of GABA, i1 is the current amplitude measured in the absence of GABA, i2 is the relative current amplitude of the maximum response to GABA, pEC_50_ is the log of the effective dose for 50% effect (EC_50_), X is the log of the GABA concentrations, and p is a slope factor. Amplitude of the constitutive current was measured at the beginning and at the end of the recording session for each oocyte without GABA perfusion. The effect of Fipronil (100µM) on the constitutive current was evaluated at the end of the recording session for each oocyte (relative to the absolute zero current). The effect of Ivermectin was evaluated after 3 application of 10 μM IVM + 100 μM GABA (relative to control Imax 100 μM GABA). The effect of ChrodrimaninB was evaluated after 2 application of 2 μM ChroB + 300 μM GABA (relative to control Imax 300 μM GABA). All values were stored in Excel (Microsoft, Office 16), graphs and Statistical tests were done with GraphPad Prism 8. All averaged values are given as mean ± s.e.m (standard error of the mean). Student t-test, or Mann-Whitney Rank Sum test, when normality test failed, or ANOVA were used to assess the differences between mean values, with a statistical significance noted in the figures: * = p<0.05, ** = p<0.01, *** = p<0.001.

## Acknowledgments

This work used the platforms of the Grenoble Instruct-ERIC centre (ISBG; UAR 3518 CNRS-CEA-UGA-EMBL) within the Grenoble Partnership for Structural Biology (PSB), supported by FRISBI (ANR-10-INBS-0005-02) and GRAL, financed within the University Grenoble Alpes graduate school (Ecoles Universitaires de Recherche) CBH-EUR-GS (ANR-17-EURE-0003). The IBS-ISBG EM facility is supported by the Auvergne-Rhône-Alpes Region, the Fondation Recherche Medicale (FRM), the fonds FEDER and the GIS-Infrastructures en Biologie Sante et Agronomie (IBISA). We thank the CM01 and CM02 staff (especially Romain Linares, Pauline Juyoux) for local contact support and Guy Schoehn for establishing and managing the IBS-ISBG cryo-electron microscopy platform and for providing access, training and support.

The work was funded by the ANR (Pestipenta ANR-21-CE11-0016, Synaptic Bee ANR-20-CE34-0017-01), an Equipe FRM grant to HN, the CEA (PhD fellowship to TL).

## Author contributions

TL, CJB and DB expressed and purified the samples, TL and EZ collected cryoEM data, TL and HN performed cryoEM data and structural analysis, MPP and FD performed molecular dynamics simulations, TL, JN, MR, TC and PC performed electrophysiology experiments, TL and HN wrote the paper with inputs from all authors.

