## Supplemental Data for "Structures of the honeybee GABA_A_ RDL receptor illuminate allosteric modulation"

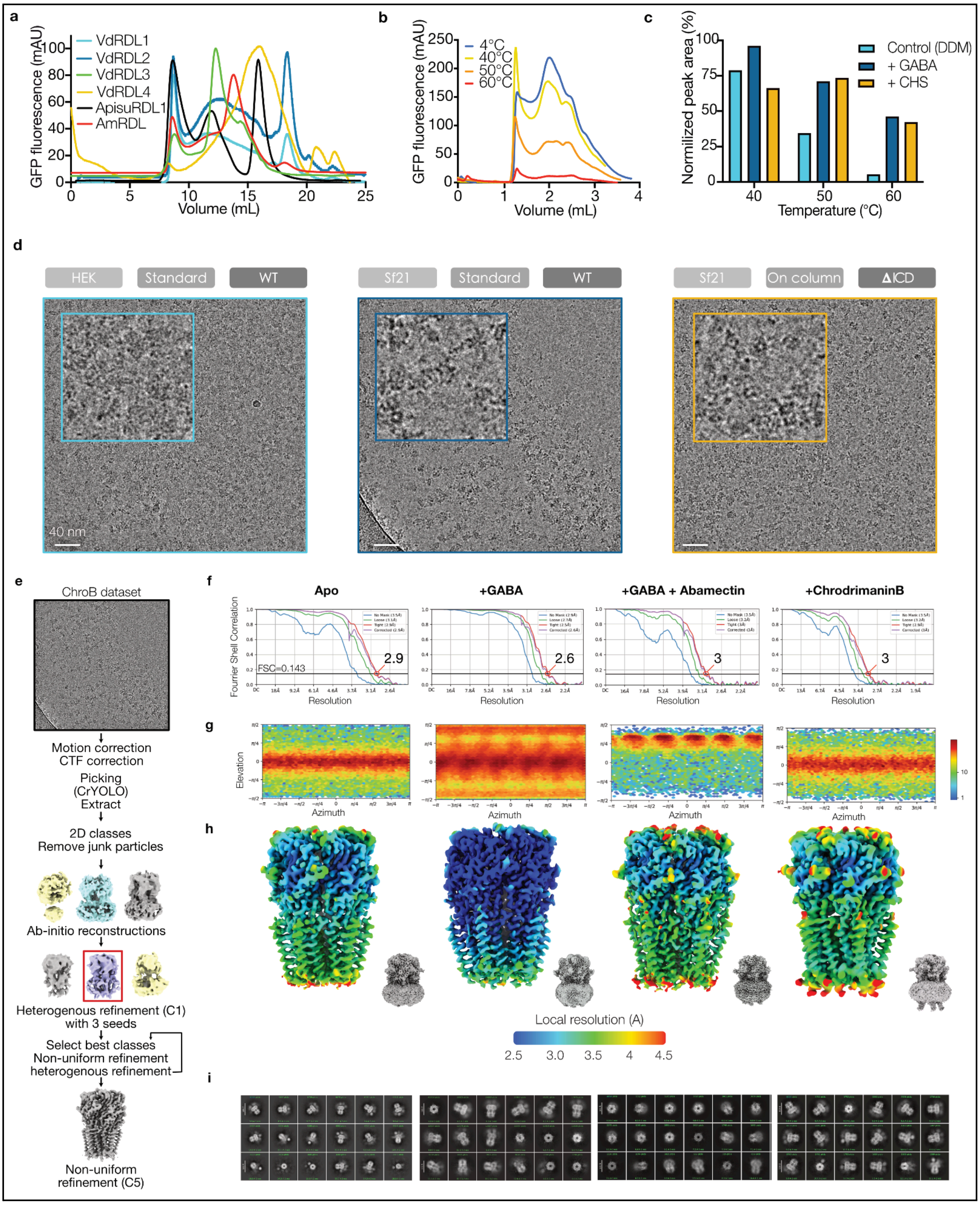


**Fig. S1: Biochemical optimization and CryoEM workflow of AmRDL receptors.**

**a.** Overlays of fluorescence size exclusion profiles of solubilized RDL receptors. Normalized to the maximum GFP fluorescence. Vd: *Varroa destructor*, Apisu: *Acyrthosiphon pisum*. **b.** Overlays of fluorescence size exclusion profiles for equal amounts of solubilized AmRDL receptors pre-heated at different temperatures to evaluate protein stability. **c.** Stability of the GFP-fused AmRDL pentamer. The sample was treated at different temperatures for 10 min in the presence of additives. The peak area corresponds to the pentamer's FSEC elution volume and was normalized to those of the samples at 4°C. Receptors were extracted with DDM or DDM:CHS (10:1 molar ratio), and incubated with 40 µM GABA. **d.** Representative micrographs of three different conditions tested. Receptors wild-type (WT) or deleted for most of the intracellular domain (334-429, ΔICD), expressed either in HEK293 or Sf21, and inserted into a lipid nanodiscs during (on-column) or after the affinity purification (standard).

**e** Schematic of a representative cryoEM data analysis workflow. **f.** Gold-standard FSC curves, line represents the 0.143 FSC threshold. **g**. Heatmap of the angular distribution of particle projections for the four reconstructions. **h.** Side views of final reconstructions. The unsharpened 3D density maps are colored according to the local resolution. **i.** 2D class averages

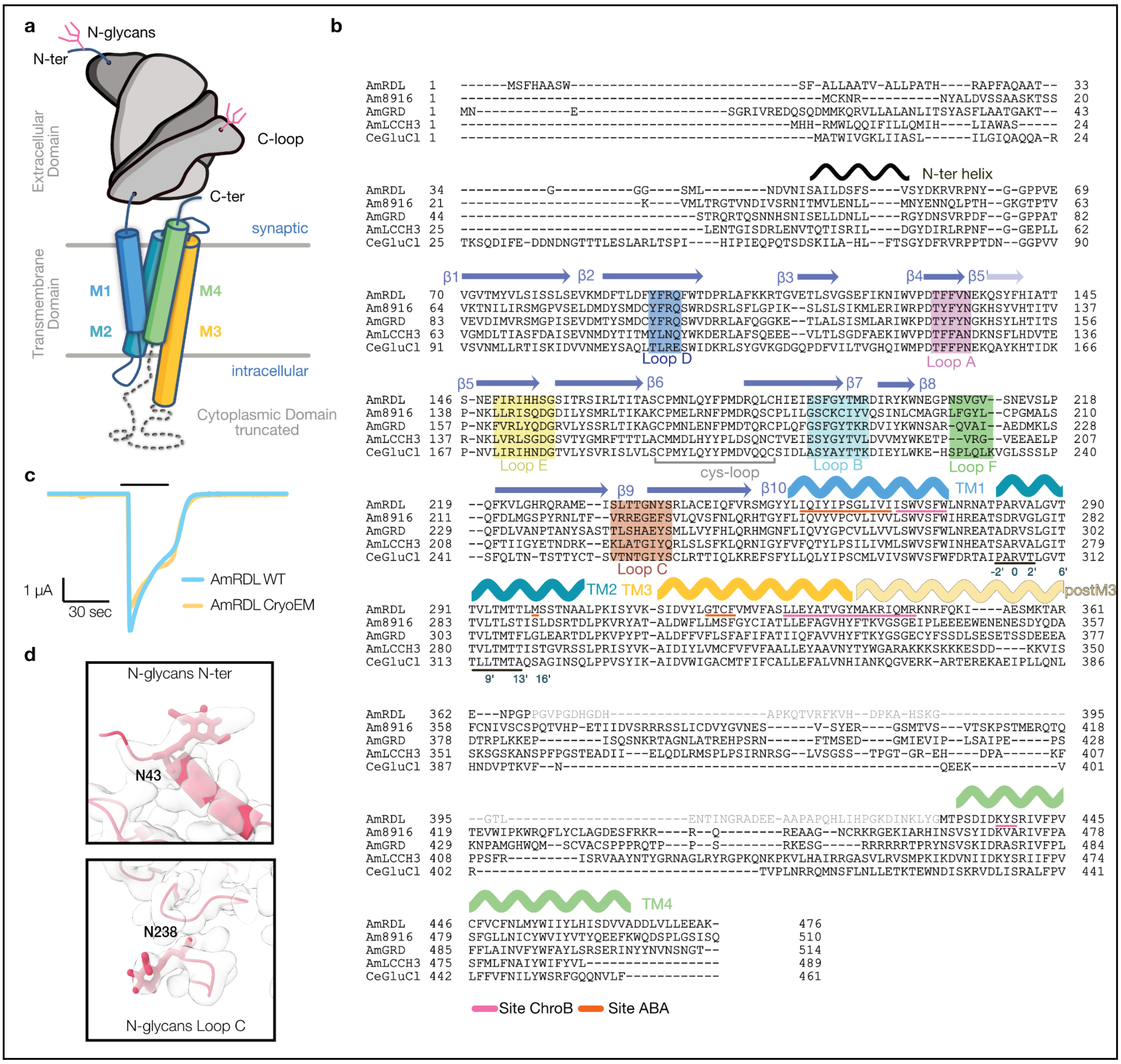


**Fig. S2 | Construct, nomenclature, sequence alignment**

**a.** Schematic of an AmRDL subunit. **b.** Multiple sequence alignment of *Apis mellifera* RDL, 8916, GRD, LCCH3 and *C. elegans* GluCl subunits. Important residues and elements of structures are highlighted. Residues highlighted in orange and pink are those belonging to the ABA and ChroB site, respectively. The gray domain corresponds to the truncated ICD domain replaced by a GFP **c.** 500 µM GABA-elicited currents in the wild-type receptor and in the construct engineered for cryoEM. **d.** Densities corresponding to N-glycosylations.

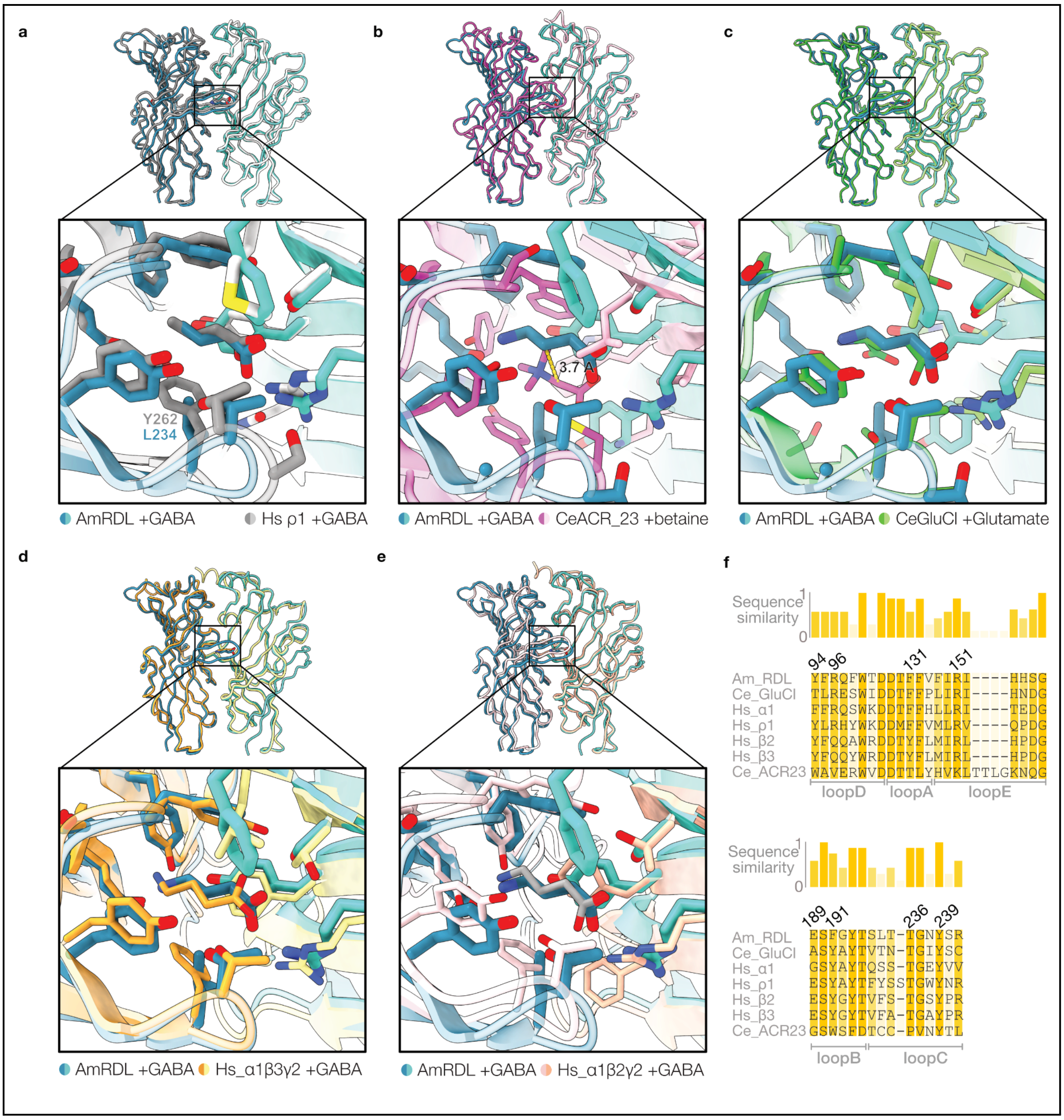


**Fig. S3 | Orthosteric binding site: comparison with homologous receptors**

**a-e.** Overlays of the GABA-bound AmRDL receptor with the GABA-bound human ρ1 (8OP9), CeACR23 betaine-bound receptor (8ZFM), the CeGluCl glutamate-bound receptor (3RIF), and the GABA-bound human α1β3γ2 (6HUJ) and α1β2γ2 (8VRN) receptor. Each panel shows a view of two neighboring ECDs superimposed on the (-) subunit, with a close up view of the binding pocket. Interacting residues are shown as sticks. **f.** Sequences alignment of the loops forming the orthosteric site. Relative similarity is indicated by color.

| Site | Mutants | GABA | | | IVM | | | ChroB | | |
|---|---|---|---|---|---|---|---|---|---|---|
| | | EC50 (μM) | SEM | n | Effect | SEM | n | Effect | SEM | n |
| | WT | 10.92 | 1.54 | 9 | 0.21 | 0.11 | 5 | 0.03 | 0.02 | 3 |
| GABA | Y239A | NR | | - | - | - | - | - | - | - |
| GABA | R96N | NR | | - | - | - | - | - | - | - |
| GABA | R151A | 15650.6 | 4373.29 | 9 | - | - | - | - | - | - |
| GABA | E189G | NR | | - | - | - | - | - | - | - |
| GABA | F191A | 1064.28 | 866.41 | 9 | - | - | - | - | - | - |
| IVM | I257F | 5.18 | 138 | 5 | 0.25 | 13 | 5 | - | - | - |
| IVM | M299S | 9.68 | 8.97 | 8 | 0.75 | 0.12 | 4 | - | - | - |
| IVM | I261F | 1.64 | 0.96 | 3 | 0.25 | 0.06 | 5 | - | - | - |
| IVM | G320F | 496.15 | 61.96 | 5 | 0.9 | 0.08 | 12 | - | - | - |
| ChroB | Y338L | 8.8 | 1.68 | 8 | - | - | - | 0.97 | 0.02 | 8 |
| ChroB | Y338A | 2530 | 903 | 7 | - | - | - | 10.49 | 3.38 | 4 |
| ChroB | W275L | 18.31 | 3.34 | 8 | - | - | - | 0.9 | 0.03 | 5 |
| ChroB | T335L | 2.4 | 0.47 | 8 | - | - | - | 0.88 | 0.01 | 8 |
| ChroB | W271Q | 3.08 | 0.87 | 6 | - | - | - | 0.92 | 0.04 | 7 |
| ChroB | A334L | 11.77 | 5.21 | 5 | - | - | - | 1.35 | 0.09 | 4 |
| | GRD-LCCH3 | 1.78 | 0.15 | 5 | 0.23 | 0.04 | 3 | 0.91 | 0.01 | 3 |

**Table S1 | Dose-response data and antagonist effect for AmRDL and mutants of the three sites**
For IVM and ChroB effect, the value represents the ratio of $I_{300uM\ GABA}$ with the inhibitor (after 2 or 3 applications for ChroB and IVM respectively) to the control $I_{300uM\ GABA}$. Data obtained from TEVC experiment in *Xenopus* oocytes. NR: not responsive.

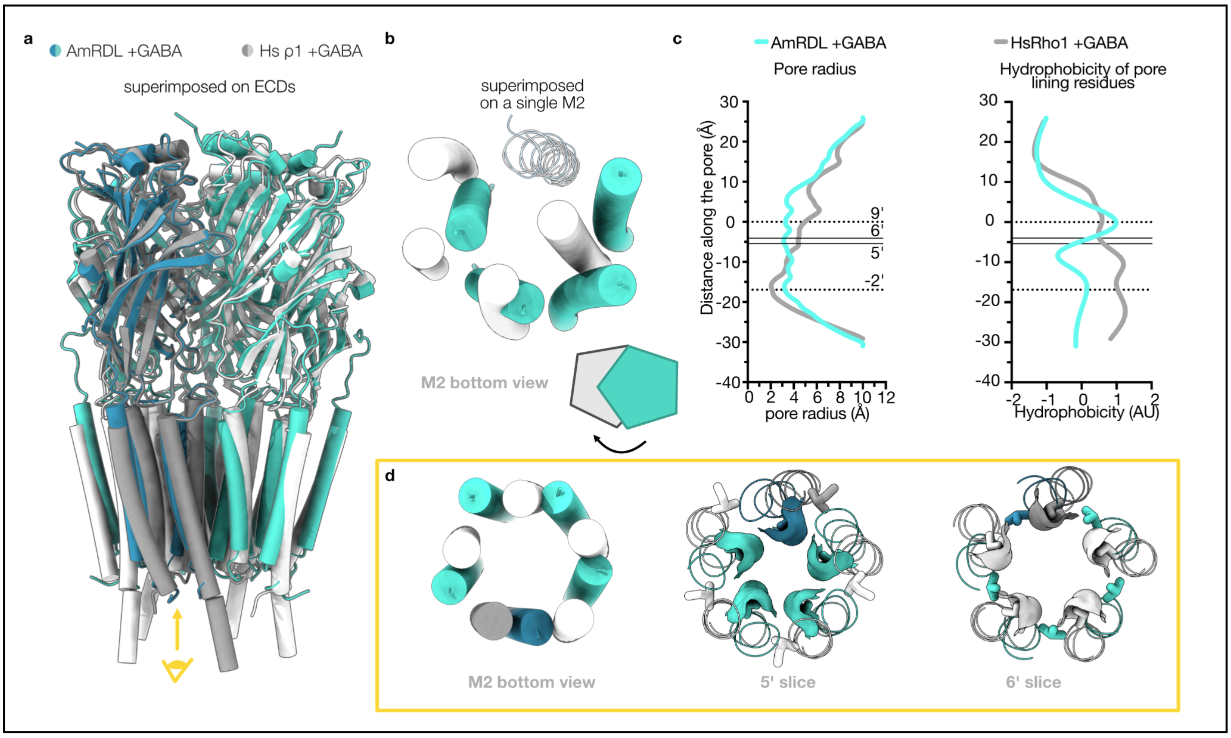


**Fig. S4 | GABA-bound conformation: comparison with the human ρ1 receptor**

**a.** Overlay of the GABA-bound AmRDL receptor with the GABA-bound human ρ1 receptor. The superimposition on the ECDs shows that ECDs adopt quite similar conformations, while there are more differences in the TMDs. The position of helices M1, M3 and M4 is still relatively similar, but large deviations are observed for M2s. **b**. When a single M2 from each structure are superimposed, the difference in the organization of the M2 bundle is striking, and the neighbors of the superimposed helix are largely offset. **c.** Pore radius and hydrophobicity of pore residues comparison. **d.** Superimposition of the M2 helical bundle also permits to see the difference of organization, when looking at the whole M2s (left) or with slabs centered on positions 5' and 6'. Note the difference in solvent-accessible surfaces: they are big for 5' for the AmRDL receptor and absent in 6' and vice versa for the ρ1 receptor.

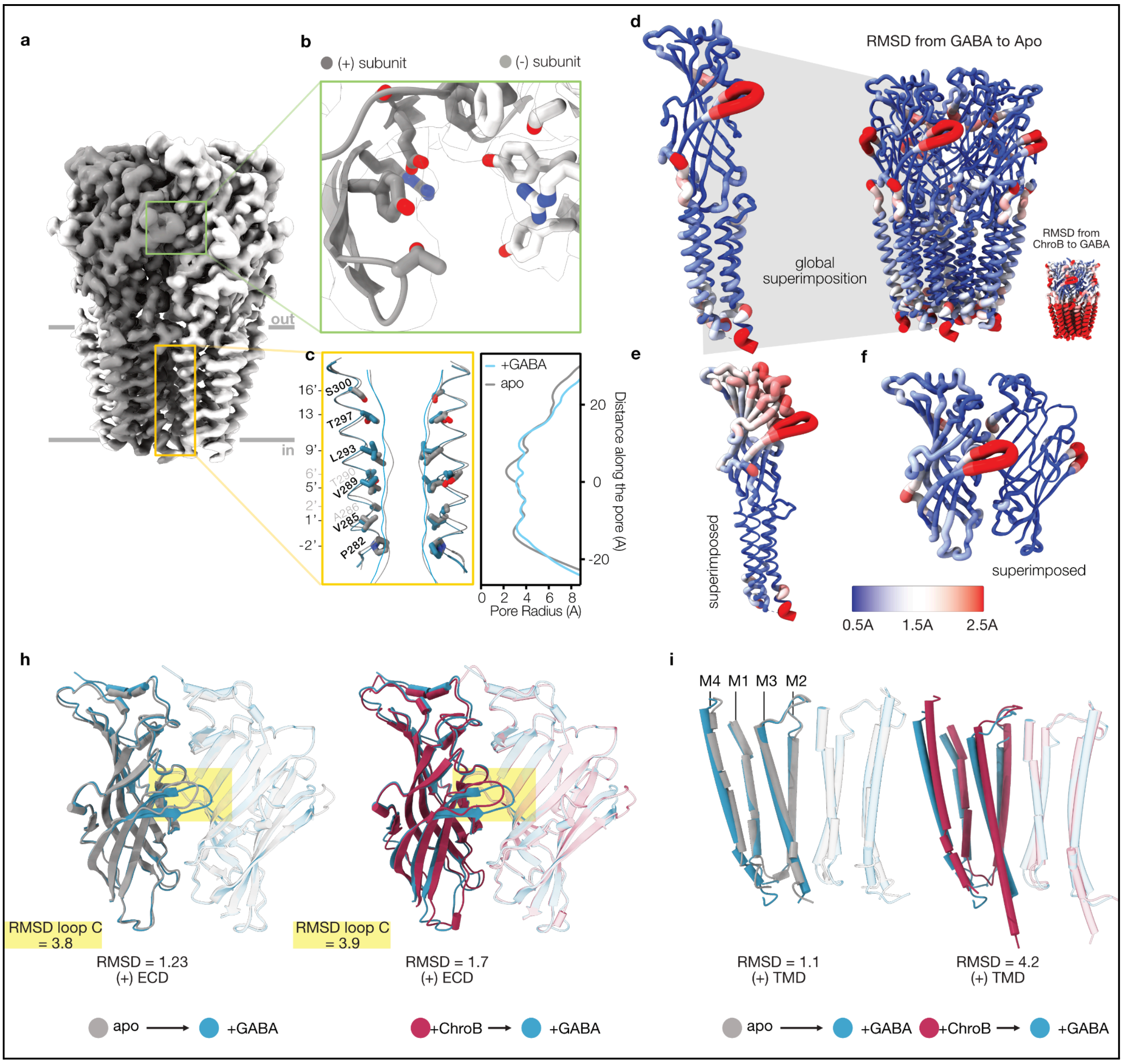


**Fig. S5| The apo AmRDL structure resembles the GABA-bound structure more than the others**

**a,** Side view of the apo-AmRDL receptor unsharpened CryoEM map. **b,** Close-up of the empty orthosteric site. **c,** Ion permeation pathway in apo-receptor. For clarity, only one M2 helix is shown as ribbons, and side chains facing the pore are shown as sticks and labeled by prime notation on the left. The accessible pathway (shown in gray line) was calculated using CHAP, and the pore profile is shown on the right. **d-f,** Structure of apo-receptor viewed from the membrane plane. The backbone color and thickness represent the r.m.s.d. relative to the GABA-bound structure. (d) global superimposition, inset: zoomed view of a single subunit. Is also represented the ChroB-bound structure relative to the GABA-bound structure (bottom right) (e) View of one subunit, superimposition on TMD. (f) View of two adjacent ECDs, superimposition of the complementary one. **h**, Overlay of two ECDs (superimposed on the complementary ECD) of the apo- and GABA-bound or ChroB-bound receptor structures, r.m.s.d are noted. **i,** Same representation as (e) for TMDs.

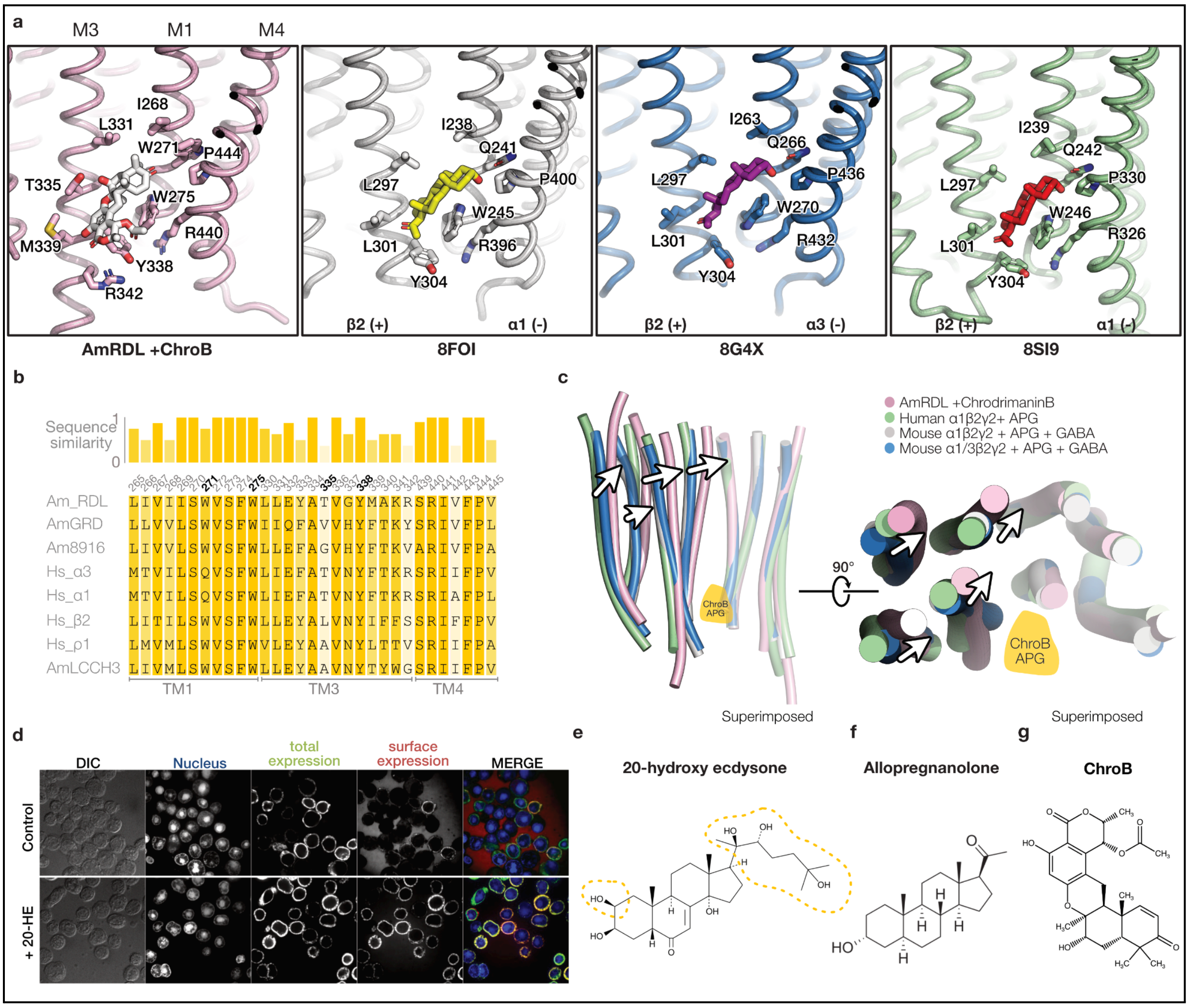


**Fig. S6 | Comparison of the ChroB and the neurosteroid PAM sites**

**a,** Side by side view of the ChroB-bound receptor, the allopregnanolone-bound α1β2γ2 and α1α3β2γ2 human receptors (PDB 8FOI and 8G5L) from left to right. Interacting residues are indicated and shown as sticks. Note the similarity in the binding pocket with strikingly conserved elements (Y338, W275, P444, R440 in RDL numbering). **b,** Sequence alignment of the neurosteroid binding sites of Human and Honeybee $GABA_A$ receptors, AmRDL residues that form the binding pocket are identified with their number. **c,** Overlays of two adjacent TMDs, from the same structures as in a, superimposed on the complementary TMD. Arrows indicate the direction of the quaternary reorganization, from the desensitized-like conformation of the neurosteroid-bound human structures to the inhibited one of the ChroB-bound receptor. **d,** Confocal images of AmRDL receptor expression in Sf21 cells in presence or absence of 20-hydroxy ecdysone (20-HE). **e-g,** Chemical structure of (e) 20-HE, (f) allopregnanolone, (g) Chrodrimanin B.

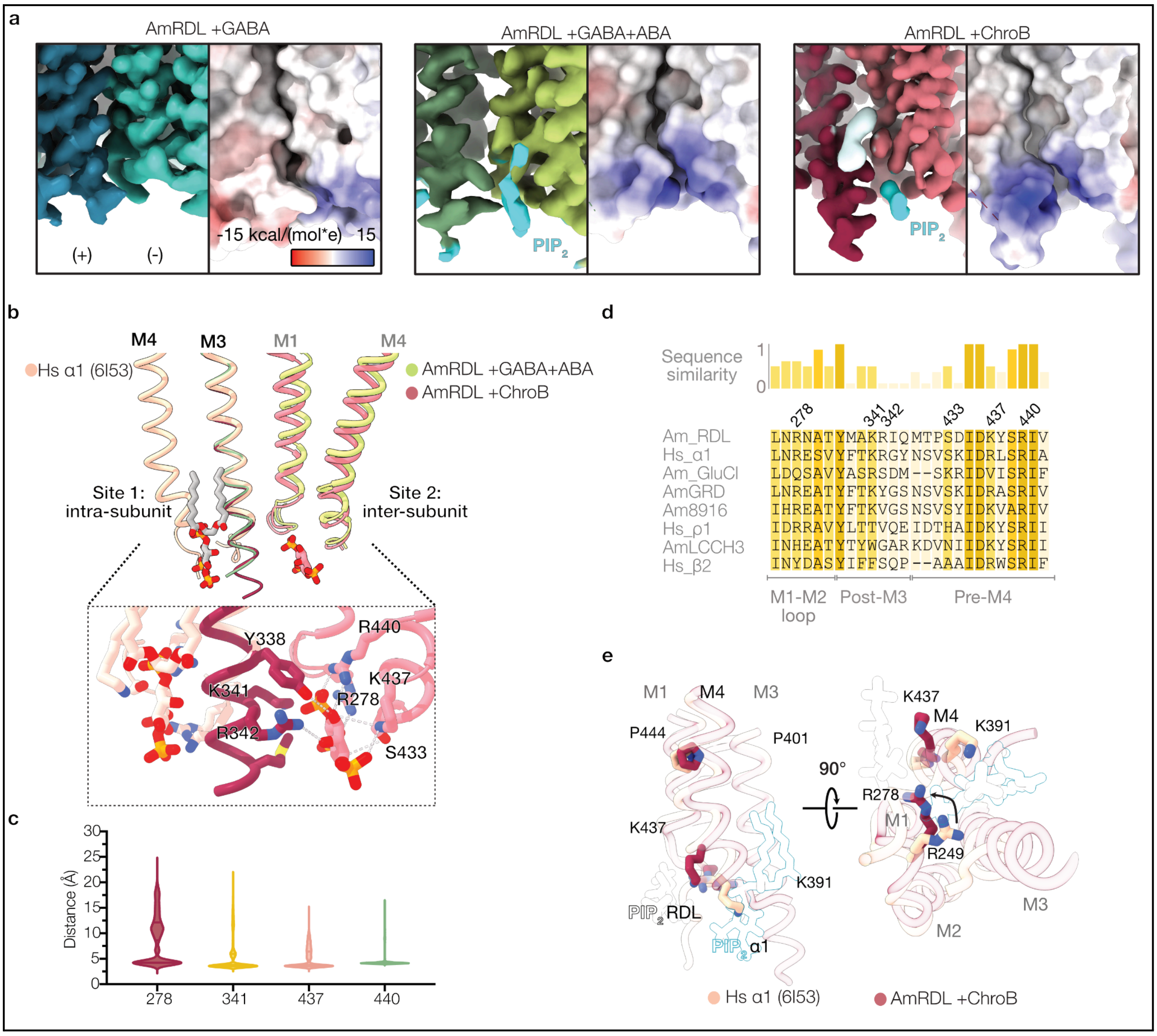


**Fig. S7 | $PIP_2$ binds to a conformation-dependant site, close to the cytoplasm and different from that of mammalian $GABA_A$ receptors**

**a**, CryoEM map (left) and electrostatic surface (right) for 3 datasets of the present work, the $PIP_2$ site is present in the ChroB and GABA+ABA conditions. In the GABA-bound condition, we note that the M1-M2 loop is not well resolved, the post-M3 is disordered from the residue 340 onwards. **b**, Overlay of $PIP_2$ site of the human α1β3γ2 receptor (camel,6I53), ChroB-bound (carmine) and GABA-ABA-bound (green) AmRDL receptors revealing two different $PIP_2$ site. Shift of the register of charged residues from the post-M3 (arrows) leads to an intersubunit site for the RDL receptor and an intrasubunit one for the human receptor. Close up of both $PIP_2$ binding sites. **c**. Distances between $PIP_2$ and binding pocket residues during MD simulations of the ChroB-bound receptor. **d**, Sequence alignment of the $PIP_2$ binding site of Human and Honeybee $GABA_A$ receptor, AmRDL residues that form H-bonds/salt-bridge with $PIP_2$ are identified. **e,** Overlays of one subunit of the human α1β3γ2 receptor (camel,6I53, $PIP_2$ underlined in cyan) and of the ChroB-bound AmRDL receptor (carmine, $PIP_2$ underlined in black). See the inversion of the site, through Arginine and lysine re-orientation.

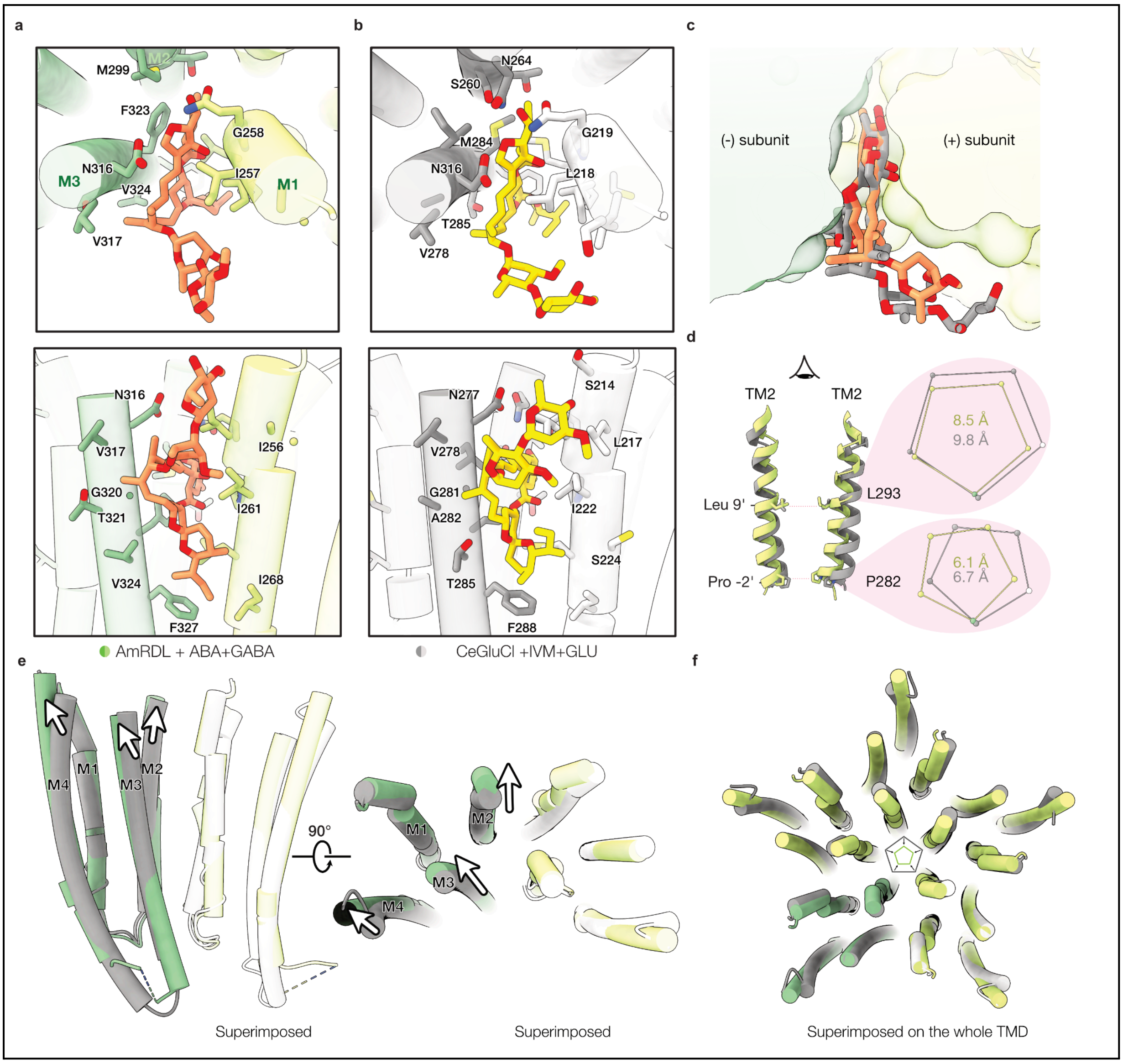


**Fig. S8 | Comparison of the GABA+ABA-bound and the GluCl Glu+IVM-bound conformations**

**a,** Views of the abamectin binding site of the GABA-ABA-bound AmRDL receptor (green) from the top (top panel) and from the side (bottom panel). Interacting residues are indicated and shown as sticks. **b,** The same representations for the ivermectin-bound CeGluCl receptor (PDB 3RIF, grey). **c,** Overlay of abamectin and ivermectin in their cavity. Note that ABA is positioned slightly deeper than IVM. **d,** Overlay of M2 helices and pentagon of Ca distances: top, Leucine 9'; bottom, Proline -2'. **e**, TMDs overlays of both receptors superimposed on the complementary TMD, with arrows indicating the direction of quaternary reorganization from the desensitized-like conformation of the IVM-bound GluCl receptor to the inhibited conformation of the GABA-ABA-bound RDL receptor. **f,** Overlays of the five TMDs viewed from the top, superimposed on the entire TMD, with arrows in the pentagon indicating the direction of TMD reorganization.

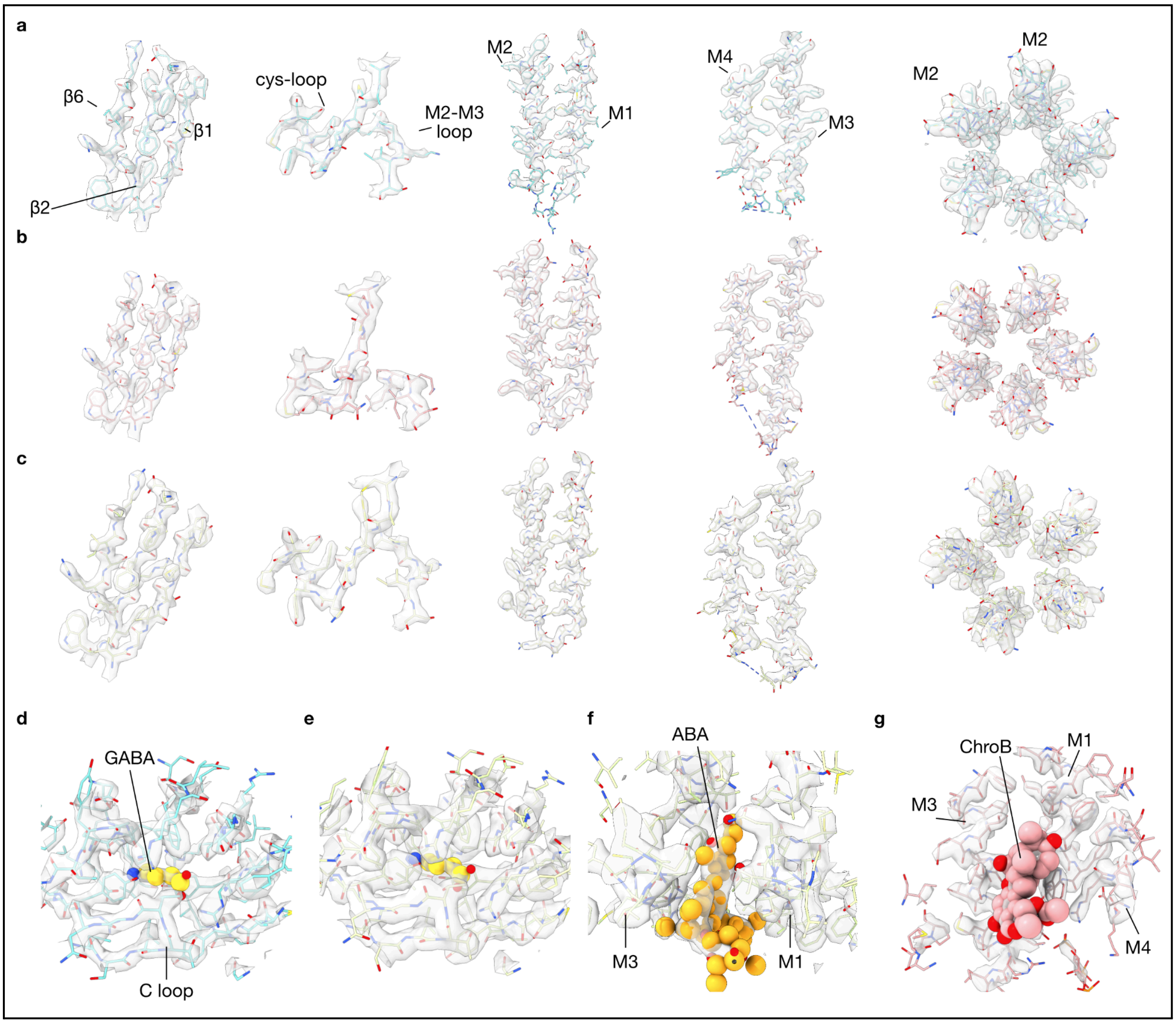


**Fig. S9 | Representative densities of the three reconstructions of AmRDL**

**a-c,** Densities are represented in transparent volume with structures of the GABA (a, cyan), ABA+GABA (b, green) and ChroB-bound (c, pink) AmRDL receptor.

**d-g**, Densities of ligand binding site. GABA binding site of GABA (d) and ABA+GABA-bound (e) receptor. ABA binding site (f) and ChroB binding site (g).

**CryoEM data collection parameters**

| | Apo receptor | GABA-bound | GABA+ABA-bound | ChroB-bound |
|---|---|---|---|---|
| **PDB ID** | XXX | XXXX | AAAA | AAAA |
| **EMDB ID** | EMD-XXXXX | EMD-XXXXX | EMD-XXXXX | EMD-XXXXX |
| Microscope | Glacios | Titan Krios | Titan Krios | Titan Krios |
| Detector | Gatan K2 | Falcon 4i | Falcon 4i | Gatan K3 |
| Magnification | 36,000 | 130,000 | 130,000 | 130,000 |
| Voltage (kV) | 200 | 300 | 300 | 300 |
| Electron exposure (e–/Å$^2$) | 42 | 40 | 40 | 41 |
| Defocus range (μm) | -0.7 to -2.1 | -0.7 to -2.1 | -0.7 to -2.1 | -0.7 to -2.1 |
| Pixel size (Å) | 1.145 | 0.94 | 0.94 | 0.839 |
| Symmetry imposed | C5 | C5 | C5 | C5 |
| Initial particle images (No.) | 1,056,384 | 2,154,358 | 1,102,000 | 1,191,620 |
| Final particle images (No.) | 54,629 | 290,250 | 30,111 | 33,309 |
| Map resolution (Å) | 2.9 | 2.56 | 3.2 | 2.96 |
| FSC threshold | 0.143 | 0.143 | 0.143 | 0.143 |
| Initial model used | Alphafold | Alphafold | Alphafold | Alphafold |

**Refinement and validation statistics**

| | Apo receptor | GABA-bound | GABA+ABA-bound | ChroB-bound |
|---|---|---|---|---|
| **Model composition** | | | | |
| Non-H protein atoms | 13,650 | 13,680 | 14470 | 14645 |
| Proteins residue | 1,690 | 1,715 | 1,745 | 1765 |
| Glycans | 5 | 10 | 10 | 10 |
| Ligands | 0 | 5 | 10 | 5 |
| Lipids | 0 | 0 | 0 | 5 |
| **B-factor (A2)** | | | | |
| (min/max/mean) | | | | |
| protein | 47/191/105 | 6/178/92 | 52/192/107 | 92/248/145 |
| ligand | 131/143/136 | 50/127/102 | 72/143/117 | 159/282/199 |
| **Ramachandrans plot** | | | | |
| Favored (%) | 97.6 | 99.09 | 98.26 | 99.14 |
| Allowed (%) | 2.4 | 0.91 | 1.74 | 0.86 |
| Disallowed (%) | 0.0 | 0.0 | 0.0 | 0.0 |
| **Validation** | | | | |
| Molprobity score | 1.52 | 1.66 | 1.49 | 1.6 |
| Clashscore | 8.03 | 14.10 | 9.12 | 12.36 |
| Poor rotamers (%) | 0.65 | 0.72 | 0.38 | 0.62 |

**Table S2 | Data collection parameters and refinement**
Table summarizing cryoEM data collection and processing statistics for the 4 datasets in this study, and cryo-EM data refinement and validation statistics for the 4 maps and models.
